# Learning a Continuous Progression Trajectory of Amyloid in Alzheimer disease

**DOI:** 10.64898/2026.02.03.703568

**Authors:** Mingzhao Tong, Fahad Mehfooz, Shu Zhang, Yipei Wang, Shiaofen Fang, Andrew J. Saykin, Xiaoqian Wang, Jingwen Yan, the Alzheimer’s Disease Neuroimaging Initiative

## Abstract

**BACKGROUND:** Understanding of early Alzheimer progression is critical for timely diagnosis and treatment evaluation, but traditional diagnostic groups often lack sensitivity to subtle early-stage changes.

**METHODS:** We developed SLOPE, an unsupervised dimensionality reduction method that models the amyloid progression in AD on a continuous scale which preserves the temporal sequence of follow-up visits. Applied to longitudinal amyloid PET data, SLOPE generated a two-dimensional trajectory capturing global amyloid accumulation across the AD continuum.

**RESULTS:** SLOPE-derived pseudotime scores better preserved temporal consistency across diagnostic groups and longitudinal follow-up visits and can be generalized to held-out subjects. The learned trajectory revealed biologically consistent amyloid spreading patterns and greater sensitivity to early progression than global amyloid SUVR.

**DISCUSSION:** SLOPE provides a continuous staging of amyloid pathology that complements global amyloid measures by capturing early localized progression.

## 1. Background

Alzheimer’s disease (AD) is a progressive, irreversible neurodegenerative disorder characterized by amyloid-β plaque accumulation. Clinically, AD develops over decades, beginning with subtle cognitive changes that gradually progress to impairments in memory and daily functioning ^1^. Despite a few FDA-approved drugs, they offer only symptomatic relief and no definitive cure exists currently ^2^. Consequently, accurately modeling disease progression and detecting subtle early-stage changes are critical for timely diagnosis and assessment of treatment effectiveness. However, the extended disease course limits longitudinal observations, as most cohorts capture only a few snap shots of each individual’s trajectory, hindering reconstruction of full temporal dynamics and identification of early pathological changes.

Traditional analyses often categorize individuals into coarse diagnostic groups, such as cognitively normal (CN), mild cognitive impairment (MCI), and Alzheimer’s disease (AD) ^3^. While convenient, these groupings fail to capture the continuous nature of AD progression, particularly in early stages when individuals may be labeled as cognitively normal or amyloid-negative despite ongoing pathology ^4^. This oversimplification masks subtle but clinically meaningful changes and limits understanding of early pathological changes critical for timely diagnosis and intervention.

In recent years, pseudotime-based approaches have emerged as effective tools for modeling disease progression trajectories by inferring latent temporal ordering from high-dimensional biomarker data in the absence of explicit time stamps. Originally developed to study cell differentiation in single-cell data, pseudotime analysis has successfully captured transitions between cell states and associated molecular changes ^5-7^. More recently, these methods have been applied to gene expression and brain imaging data to reconstruct continuous sequences of pathological changes across individuals at different disease stages ^8-10^.

Despite their promise, pseudotime methods have notable limitations. They are often non-parametric and primarily used for exploratory analysis, with limited ability to generalize to unseen patients. In addition, most approaches are applied to cross-sectional data and do not explicitly incorporate temporal context from longitudinal follow-up visits, which is essential for modeling disease progression. The lack of standardized benchmarks for evaluating inferred trajectories further limits assessment of reliability and reproducibility.

To address these challenges, we present a novel Self-supervised Longitudinal Progression Embedding (SLOPE) method for modeling amyloid progression in Alzheimer’s disease. SLOPE integrates high-dimensional biomarker data with temporal information from longitudinal follow-up visits in a parametric framework, producing a visually intuitive two-dimensional progression trajectory that supports individualized staging for both training and unseen subjects. We further introduce two metrics to evaluate trajectory robustness. Given that amyloid accumulation is considered one of the earliest pathological features of AD ^11^, we applied the SLOPE method to longitudinal amyloid PET imaging data from the Alzheimer’s Disease Neuroimaging Initiative. Compared with state-of-the-art methods, SLOPE more accurately preserved temporal ordering of follow-up visits and showed greater sensitivity to early regional amyloid accumulation than global amyloid SUVR.

## 2. Methods

### 2.1 Dataset

Longitudinal Amyloid PET data was downloaded from the Alzheimer’s Disease Neuroimaging Initiative (ADNI) database, which was processed by the UC Berkeley group and had already undergone pre-processing steps, providing direct regional amyloid standardized uptake value ratios (SUVRs) for each brain region. Details of image preprocessing and SUVR estimation are available in ADNI documentation.

This dataset includes florbetapir amyloid PET data from 961 individuals across 2,023 visits; data from other amyloid PET tracers were excluded to ensure consistent SUVR scaling. Each visit was assigned to one of four diagnostic groups: cognitively normal (CN), early MCI (EMCI), late MCI (LMCI), or Alzheimer’s disease (AD). Amyloid burden was quantified using SUVRs from 68 cortical regions, normalized by the ADNI-provided composite reference region COMPOSITE REF SUVR. This normalization step helps control for nonspecific binding and individual variability, ensuring that measurements accurately reflect true amyloid deposition. Subcortical regions were excluded from the amyloid analysis, as their amyloid burden is less specific and not closely associated with AD. Imaging measures were pre-adjusted for age, sex, and education using weights derived from baseline CN participants. All imaging features were standardized to have a mean of zero and a standard deviation of one for subsequent analysis. This center-scaling of amyloid SUVR values was applied solely to improve the training stability of proposed SLOPE model and does not alter the biological interpretation of amyloid positivity, as the model learns relative regional amyloid patterns rather than relying on absolute SUVR thresholds.

The cohort included 395 individuals with baseline-only visits and 526 with longitudinal follow-up, capturing transitions across disease stages (Fig. 1a). Among those with only baseline data, there are 70 CNs, 102 EMCIs, 83 LMCIs, and 140 AD patients. All individuals, regardless of whether they had follow-up visits, were included in the trajectory learning and evaluation.

**Figure 1:**
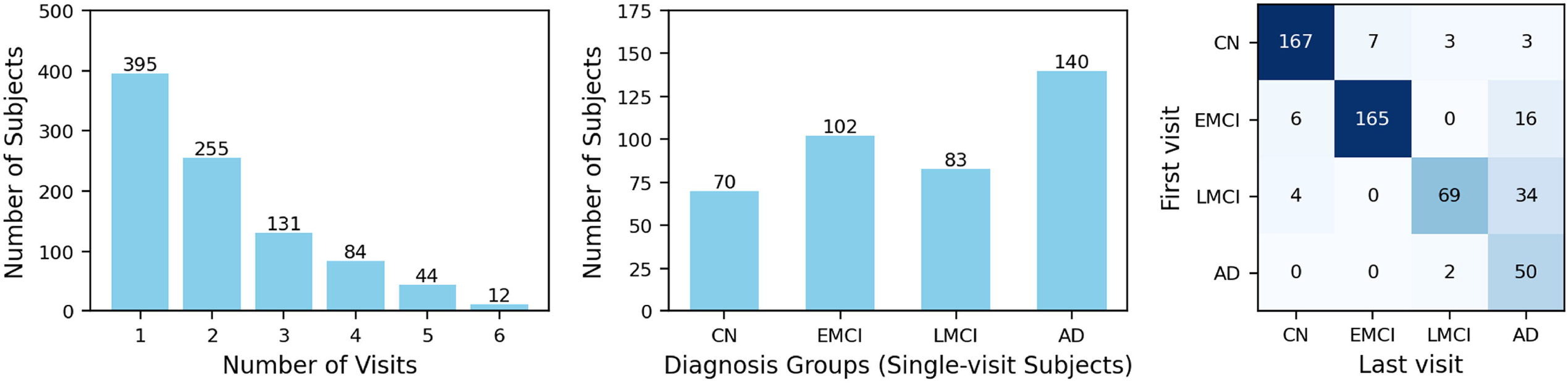
(a) Distribution of ADNI participants with amyloid-PET data. Left: Number of follow-up visits among all participants. Middle: Diagnosis distribution of participants with only baseline visit. Right: diagnosis distribution of participants with multiple follow-up visits. Heatmap represents the diagnosis change from first to last visit. (b) Overview of the proposed SLOPE model. It will generate a low-dimensional embedding that preserves the temporal order of each individual’s follow-up visits. This embedding will be further reduced into 2D via UMAP, which is expected to preserve the temporal order of follow-up visits learned from SLOPE and form a 2D disease progression trajectory, reflecting the continuous process of amyloid accumulation in the brain (c).

Detailed demographic information is in Table. 1.

### 2.2 Self-supervised Longitudinal Progression Embedding (SLOPE)

We used a two-step approach to derive a compact two-dimensional disease progression trajectory with temporally ordered follow-up visits. The proposed SLOPE model first reduces regional amyloid SUVR dimensionality while preserving the temporal order of follow-up visits (Fig. 1b), and the resulting embeddings are mapped to two dimensions for visualization of progression (Fig. 1c). The objective is for follow-up visits to align with a common progression direction with minimal reversals.

SLOPE is an autoencoder-based neural network trained in a fully self-supervised manner to learn low-dimensional embeddings of regional amyloid-β levels without diagnostic labels. The model architecture consists of two main components:

- **Encoder** *g*_*ϕ*_: maps the amyloid SUVR values of 68 brain regions for subject *i* at visit *k*, denoted as **x**_*i,k*_, into a low-dimensional embedding **z**_*i,k*_.
- **Decoder** 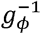 : reconstructs the original amyloid SUVR vector from the embedding.

The autoencoder is trained by minimizing a reconstruction loss that measures the discrepancy between the observed amyloid profile and the decoder’s output:

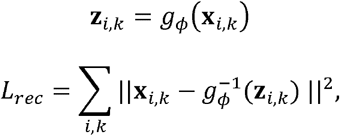

To model longitudinal follow-up data, we introduce a direction loss that enforces a biologically plausible progression trajectory by encouraging non-decreasing amyloid accumulation across visits. This loss constrains each subject’s latent trajectory to follow a consistent progression direction, reflecting the assumption that amyloid burden does not spontaneously reverse in untreated Alzheimer progression. For subject *i*, the progression vector between consecutive visits *k* and *k* + 1is defined as

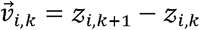

The direction loss *L*_dir_penalizes abrupt changes in progression by encouraging alignment between consecutive progression vectors via cosine similarity. Sharp directional changes reduce similarity and incur higher penalties, discouraging irregular trajectories such as those illustrated in Fig. 1c.

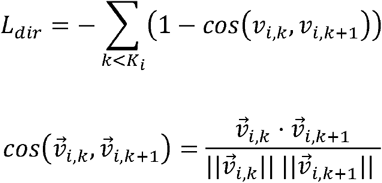

The training process of SLOPE aims to control the reconstruction loss and direction loss simultaneously, by minimizing the weighted total loss *L*_*total*_ =*λ*_1_ *L*_*rec*_ + *λ*_2_ *L*_*dir*_ Here, *λ*_1_ and *λ*_2_ are two hyperparameters that control the relative contribution of two losses. The direction loss was applied only to subjects with two or more longitudinal visits, whereas subjects with a single visit contributed exclusively to the reconstruction loss and did not influence temporal ordering constraints. Together, these losses encourage a smooth, compact embedding that captures both cross-sectional similarity and longitudinal disease progression.

### 2.3 Training and Test Data Split

The entire amyloid-PET dataset was split at the subject level to prevent any potential data leakage between the training and test sets. To preserve the distribution of diagnostic groups across both subsets, we employed stratified random sampling with a fixed random seed. This ensured that 80% of unique subjects were assigned to the training set and the remaining 20% to the test set. To ensure a fair comparison, the same data partition was applied across all competing methods. Details of hyperparameter tuning and optimal parameters can be found in supplementary text.

### 2.4 Pseudotime calculation

The low-dimensional embeddings learned from SLOPE were further transformed using a parametric UMAP model, resulting in a two-dimensional representation of the progression trajectory. Within this 2D space, we fitted a principal curve that traces the overall direction of progression trajectory. Amyloid data from each visit of training subjects was then projected onto this curve, and its relative position was computed using the SlingShot package [4]. These positions were subsequently normalized to the range [0,1], yielding a continuous measure (namely pseudotime) that reflects the global amyloid burden across the brain. Once the SLOPE and UMAP models were trained, the same procedure was applied to previously unseen test subjects. Specifically, the regional amyloid SUVR values from each visit were passed through the pre-trained SLOPE and UMAP models and then projected onto the 2D progression trajectory. This projection enabled us to assign a pseudotime value to each visit, representing its position along the full spectrum of amyloid progression.

### 2.5 Model evaluation

We evaluated SLOPE by examining how well its derived pseudotime preserves temporal ordering across diagnosis groups and follow-up visits. SLOPE was benchmarked against a traditional autoencoder and supervised models including logistic regression, elastic net, and multilayer perceptron (MLP). These supervised models were trained to discriminate AD from CN, yielding a continuous logit score for each visit, which was normalized into [0,1]. We compare these logits with pseudotime from SLOPE and the autoencoder in terms of their ability to preserve the temporal sequence across diagnosis groups and follow-up visits. Progression across diagnosis groups was assessed by comparing score distributions using t-tests.

Temporal consistency across follow-up visits was evaluated using two metrics: the violation ratio and the violation gap. The violation ratio *r* measures the proportion of consecutive visit pairs that violate the expected monotonic progression assumption, where pseudotime should remain stable or increase over time. The violation tolerance parameter τ is set to allow small reversals of pseudotime between follow-up visits to be ignored when assessing temporal ordering. A reversal exceeding a tolerance threshold was considered a violation, with lower values of *r* indicating better temporal consistency. Here, *N* is the total number of subjects, *K*_*i*_ is the number of visits for subject *i, p*_*i,k*_ is the pseudotime value for subject *i* at visit *k* and **1** [·] is the indicator function that equals 1 if the condition is true and 0 otherwise.

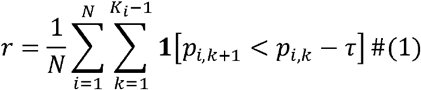

The violation gap *g* quantifies the average magnitude of such reversals, reflecting their severity rather than frequency:

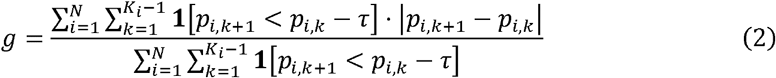

### 2.6 Comparison of SLOPE Pseudotime with Composite Amyloid SUVR

To examine whether a temporally preserved amyloid staging axis captures progression beyond global amyloid burden, we performed three analyses comparing SLOPE pseudotime with global composite amyloid SUVR. First, we assessed differences between CN and EMCI participants using both measures, performing analyses separately in the training and held-out test sets to evaluate generalization. Second, we examined the relationship between composite amyloid SUVR and SLOPE pseudotime using two-segment piecewise linear regression, allowing for nonlinear associations along the staging axis. Third, to assess whether early changes in pseudotime reflect biologically meaningful signals, we evaluated correlation between early pseudotime values and regional amyloid SUVRs across 68 cortical regions.

## 3. Results

### 3.1 Progression trajectory

The low-dimensional embeddings learned from SLOPE further went through UMAP to construct a two-dimensional progression trajectory, with all visits of training subjects ordered along a principal curve (Fig. 2a). The relative position of each visit on the principal curve was quantified as pseudotime, reflecting the global amyloid progression stage. Each visit was then assigned to a relative position along this curve, normalized to the range [0, 1], representing its pseudotime value (Fig. 2b).

**Figure 2:**
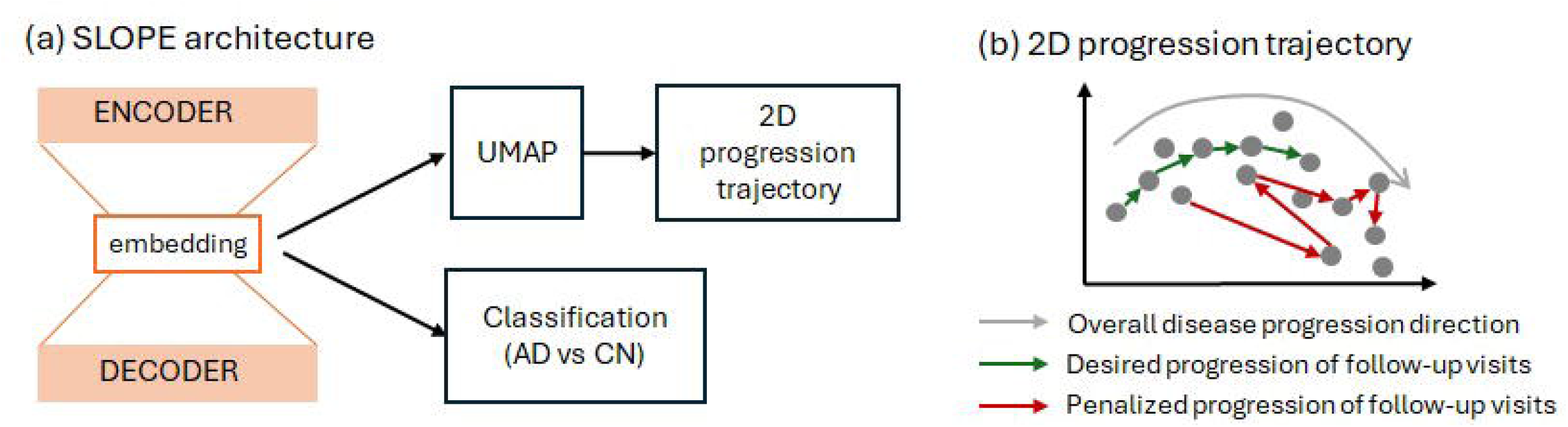
Performance comparison of SLOPE and baseline methods in preserving progressive pattern across diagnosis groups and follow-up visits. a–b: SLOPE-derived progression trajectories for training subjects, colored by diagnosis group (a) and pseudotime (b). c: Pseudotime distribution across all visits of training subjects, stratified by diagnosis group. d–f: Test subjects projected onto the SLOPE-derived progression trajectory and their pseudotime distribution. g–i: progression trajectory and pseudotime distribution derived from traditional autoencoder. j-l: Distribution of normalized logits across diagnosis groups, generated using logistic regression (j), elastic net regression (k), and a multilayer perceptron (MLP) (l), all evaluated on test subjects.

We compared the distribution of pseudotime across diagnostic groups (Fig. 2c). As expected, CN participants exhibited significantly lower pseudotime values compared to both EMCI and LMCI groups (*p* = 1.3 × 10^−17^), while AD patients showed the highest pseudotime values overall (*p* = 3 × 10^−24^)However, there was no noticeable difference between the EMCI and LMCI groups in terms of pseudotime distribution. Importantly, when the visits of held-out test subjects were projected onto the same progression trajectory, we observed a consistent pattern of group separation (Fig. 2e-g), supporting the robustness and generalizability of the learned trajectory. Similar as SLOPE, we estimated pseudotime using an autoencoder without incorporating temporal information across follow-up visits. We observed a highly consistent pattern between the training and held-out test datasets, with an incremental pseudotime distribution from CN to MCI and then AD groups.

In contrast, the three supervised models—logistic regression, elastic net, and multilayer perceptron (MLP)—were trained as multi-class classifiers. To evaluate their ability to capture the progressive nature from CN to AD, we examined the distribution of their output logits (normalized to the [0,1] range) across diagnostic groups. Unlike SLOPE and the autoencoder, the normalized logits generated from the logistic regression and elastic net models showed statistically significant differences between diagnostic groups, but lacked a clear increasing trend from CN to MCI and AD. This suggests that while these models may distinguish between groups, they do not capture the progressive nature of amyloid accumulation. Among the supervised models, only the MLP produced logits that followed a broadly increasing pattern from CN to MCI to AD, partially reflecting the amyloid accumulation along disease progression.

### 3.2 Temporal consistency of follow-up visits in progression trajectory

We further evaluated the progression trajectory by assessing how well it preserved the temporal order of longitudinal follow-up visits. For participants with multiple visits, we expect the pseudotime to either remain stable or increase over time, reflecting the irreversible nature of amyloid accumulation. To quantify this, we computed two metrics: the violation ratio and the violation gap. Both metrics were calculated across a range of tolerance thresholds, from 0.05 to 0.25 at the interval of 0.05, which allow for the exclusion of minor fluctuations in pseudotime.

Fig. 3a shows the violation ratio, which reflects the proportion of consecutive visits with pseudotime reversal exceeding a specified tolerance threshold, where the later visit has a pseudotime value lower than the earlier one. Fig. 3b presents the corresponding violation gap, which quantifies the average magnitude of those reversals. Across tolerance thresholds, the violation ratio decreased sharply as minor visit-to-visit fluctuations were progressively ignored. Under zero tolerance (τ = 0), approximately 35% of consecutive visits showed reversals in SLOPE-derived pseudotime. Allowing a small tolerance of 0.05 reduced this to ∼10%, and at τ = 0.1 only ∼4% of consecutive visits showed reversal, with an average violation gap of ∼0.12. Beyond this point, the violation rate plateaued. This pattern indicates that most reversals observed under very small τ values reflect minor visit-to-visit fluctuations rather than actual regression in disease stage. In contrast, other methods consistently exhibited both higher frequencies of reversals and larger reversal gaps across thresholds. Notably, SLOPE-derived pseudotime showed no reversals greater than 0.15, which is substantial given the total range [0,1] and further highlights its strength in preserving temporal consistency of follow-up visits.

**Figure 3:**
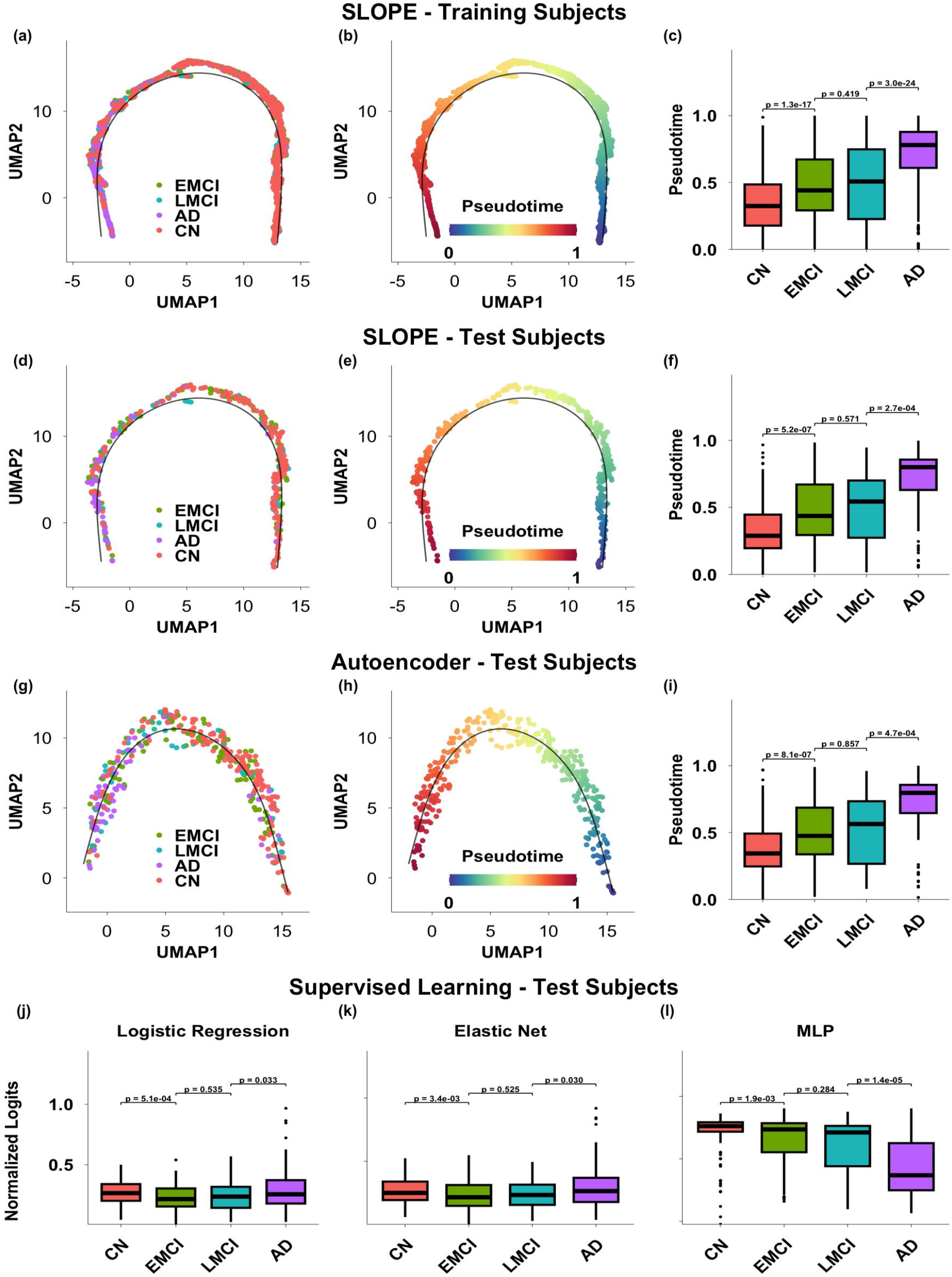
Temporal consistency across follow-up visits evaluated through violation ratio (a) and violation gaps (b). In (c) and (d) are a few example training and test subjects highlighted on the progression trajectory derived from SLOPE. Each dot is a visit, and each subject is represented as a sequence of connected follow-up visits.

Taking together, these results demonstrate the robustness of SLOPE in modeling disease progression in a temporally consistent manner, making it a strong candidate for tracking disease progression over time. In Fig. 3c-d we highlighted the follow-up visits from several example training and test subjects. This 2D progression trajectory enables direct visualization of individual-level progression dynamics, providing a practical tool for visually tracking the disease development of patients and monitoring the treatment outcomes in clinical settings.

### 3.3 Comparison with global SUVR in capturing early amyloid progression

In the training set, composite amyloid SUVR differed significantly between CN and EMCI groups (*p* = 1.29 × 10^−5^) (Fig. 4a), though this separation was much weaker than that observed with SLOPE pseudotime (*p* = 1.3 × 10^−17^) (Fig. 2c). In the held-out test set, composite amyloid SUVR did not differentiate CN and EMCI groups (Fig. 4a), whereas SLOPE pseudotime continued to show a significant separation (*p* = 5.2 × 10^−7^) (Fig. 2f), indicating improved robustness for early-stage differentiation. Segmented regression revealed a nonlinear relationship between composite amyloid SUVR and SLOPE pseudotime (Fig. 4b). In both training and test datasets, composite SUVR remained relatively stable until pseudotime reached approximately 0.36, after which it became strongly correlated with pseudotime (r ≈ 0.8). Regional analyses showed that early pseudotime variation (before 0.36) was strongly associated with amyloid accumulation in specific cortical regions, most prominently the posterior cingulate cortex (Fig. 4c) ^12,13^.

**Figure 4:**
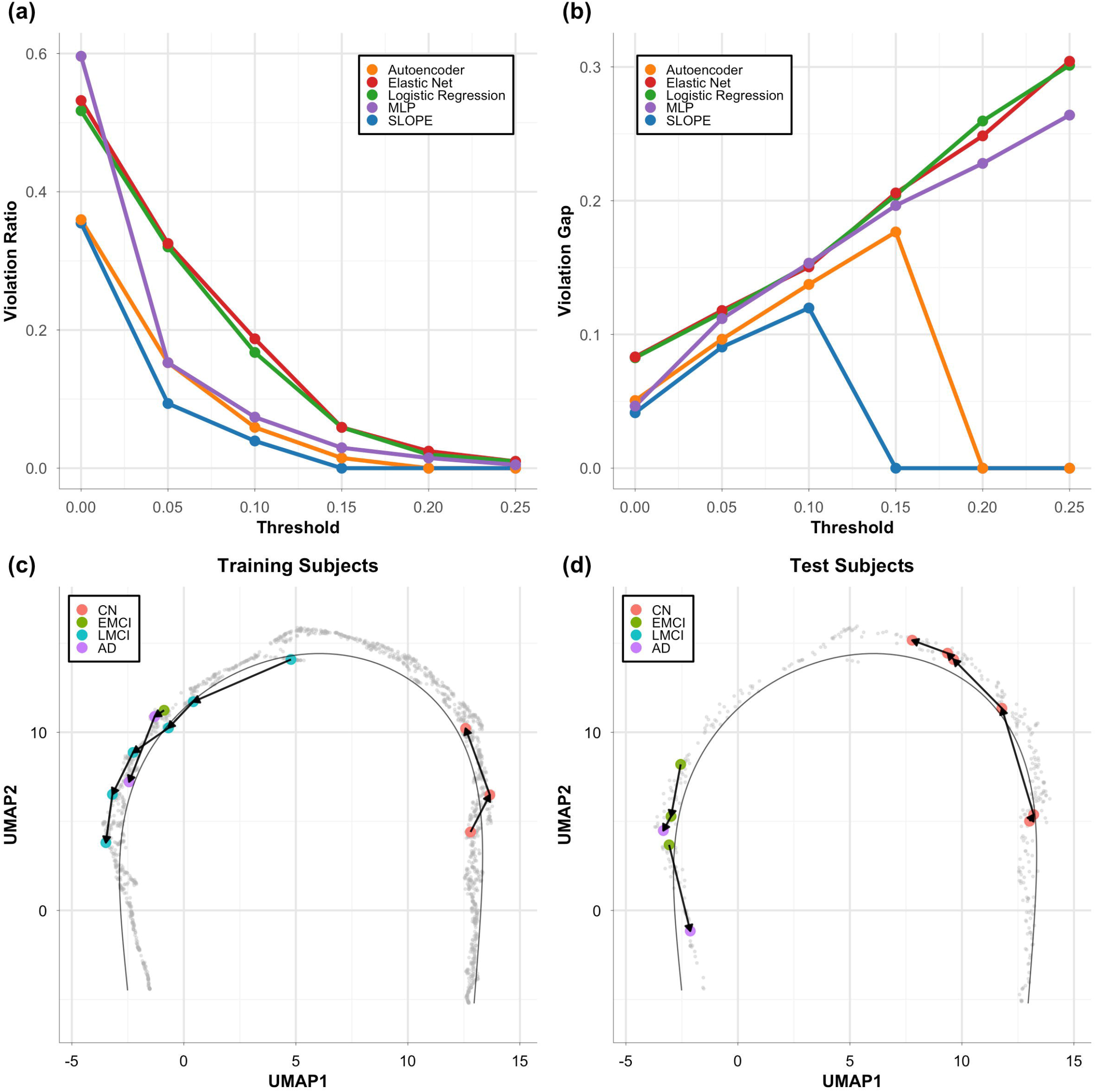
Comparison of SLOPE pseudotime and global amyloid SUVR. (a) distribution of SLOPE pseudotime and composite SUVR in CN and EMCI groups for training and test datasets. (b) Two-segments linear regression between SLOPE pseudotime and composite SUVR. (c) correlation of early pseudotime change (<0.36) with 68 regional amyloid SUVR features.

### 3.4 Brain changes along the progression trajectory

We performed a clustering analysis across all visits along the trajectory. This clustering analysis resulted in 16 clusters organized along the progression trajectory (Fig. 5a). The number of clusters was chosen for visualization purposes to provide sufficient granularity. Using Cluster 0 as the reference (representing the earliest stage), we computed the median regional SUVR values within each subsequent cluster and quantified their deviation from Cluster 0 using the following equation::

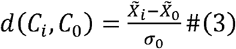

**Figure 5:**
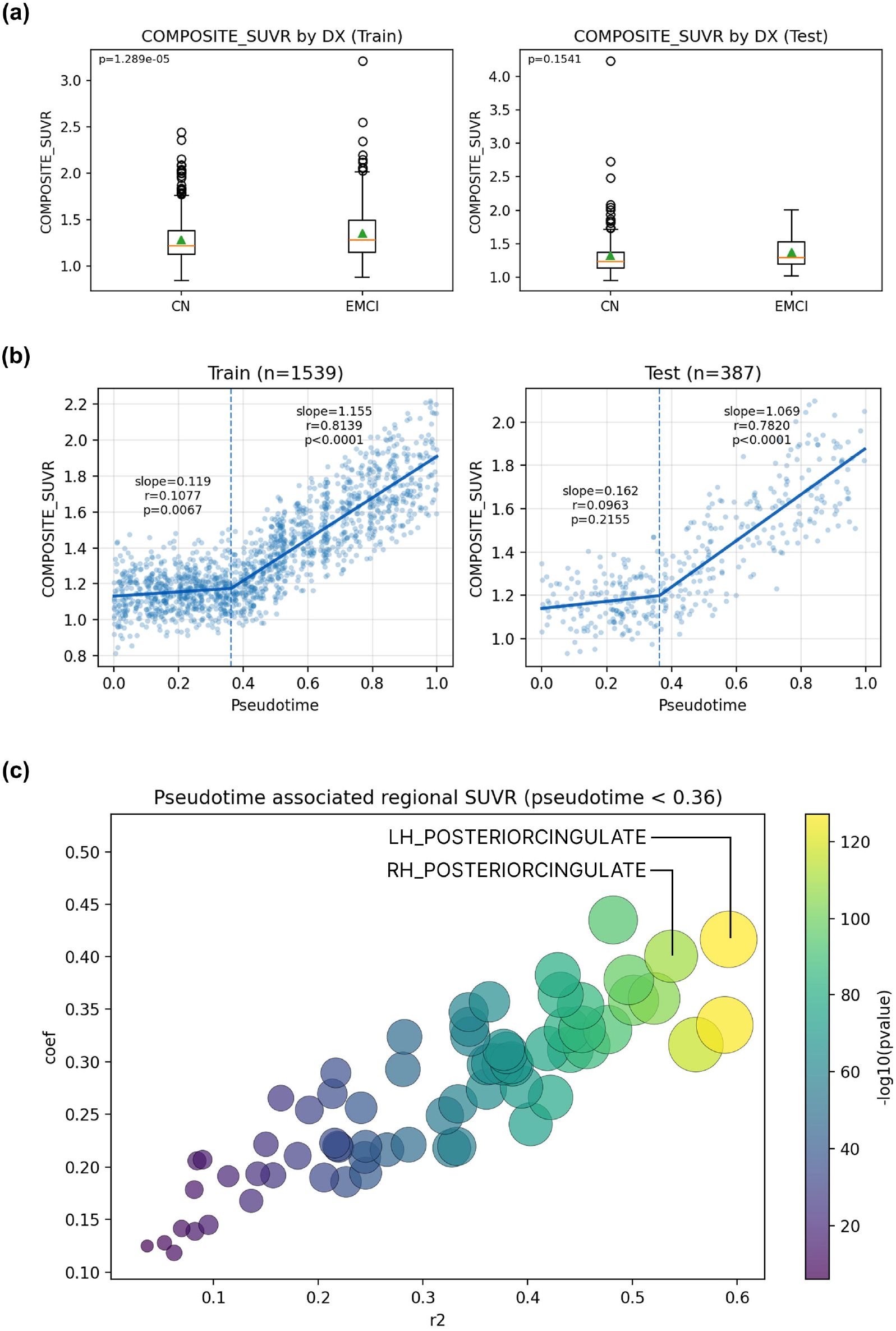
(a) Sixteen clusters were identified along the two-dimensional SLOPE-derived amyloid progression trajectory. (b) For each of the 68 bilateral cortical regions, we quantified the deviation of subsequent clusters relative to the earliest cluster (Cluster 0). (c, d) Early amyloid elevation was observed in the precuneus, posterior cingulate, superior parietal, and superior frontal cortices.

Here, for each brain region, 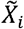 and 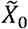 represent the median SUVR values in Cluster *i* and Cluster 0, and *σ*_0_ is the standard deviation of Cluster 0. This deviation captures how much the amyloid-β uptake levels in each brain region diverges from the baseline (Cluster 0) as the disease advances.

An initial noticeable elevation in amyloid-β uptake levels was observed in the posterior cingulate and precuneus (Fig. 5b).. Their overall amyloid uptake levels start high and start to increase rapidly from the very early stage. Both regions are core components of the default mode network (DMN), aligning with previous reports that DMN structures are particularly vulnerable to early amyloid deposition ^13^. The amyloid burden within the superior frontal cortex also demonstrated a noticeable deviation from cluster 0 in the early stage. However, upon closer examination, we found that SUVR values in the superior frontal cortex remained substantially lower than those observed in the precuneus and posterior cingulate. This pattern may reflect contributions from its anteromedial and medial subregions, which are also part of the DMN and have been suggested to be among the first sites of amyloid accumulation in the preclinical stages of AD. Another core hub of default mode network, the inferior parietal cortex, also showed signs of early elevation in amyloid uptake level, as consistently implicated in previous studies ^13^. Furthermore, the superior parietal lobe was found to undergo a relatively rapid increase in the early stage as well. But its overall amyloid uptake is substantially lower compared to early hubs such as the precuneus, posterior cingulate, and inferior parietal cortex (Fig. 5c-d). This suggests that while the superior parietal lobe is not among the first regions to accumulate amyloid, it can show fast early growth once deposition is initiated, reflecting its vulnerability as the pathology spreads outward from the core default mode network hubs. In contrast to regions within the default mode network, the precentral and postcentral cortices, together with the lateral occipital cortex, showed little deviation until approximately Cluster 9, suggesting that these areas did not experience substantial amyloid-β elevation until later stages of disease progression. This observation aligns with prior studies that have consistently identified these regions as relatively late in the amyloid accumulation sequence ^14^.

## 4. Discussion

In this work, we introduced SLOPE, a self-supervised dimensionality reduction framework for modeling amyloid progression on a continuous scale across Alzheimer’s disease development. By learning directly from longitudinal amyloid PET data without diagnostic labels, SLOPE orders all visits along a principal curve, yielding a compact two-dimensional progression trajectory that captures both global amyloid changes and the temporal ordering of follow-up visits. This trajectory preserves progression patterns across diagnostic groups as well as within individuals over time, reflecting both inter- and intra-subject disease dynamics.

Several features distinguish SLOPE from existing approaches. As a parametric model, SLOPE enables projection of unseen subjects onto the learned trajectory, supporting both retrospective analysis and prospective applications such as progression monitoring in clinical trials. Unlike traditional unsupervised clustering or supervised classification methods, SLOPE explicitly incorporates longitudinal ordering, reflecting the irreversible nature of amyloid accumulation in the untreated scenario. By preserving temporal sequence across follow-up visits, SLOPE produces a biologically meaningful and clinically interpretable disease trajectory.

The comparison between composite amyloid SUVR and SLOPE pseudotime highlights the limitations of global summary measures for early disease characterization. Composite SUVR is dominated by regions with high amyloid burden and primarily reflects mid-to-late stage pathology, making it less sensitive to subtle regional changes during early progression. In contrast, SLOPE is trained to capture localized amyloid changes for differentiation of follow-up visits even when global composite SUVR remains stable. This property further explains its improved sensitivity to early progression. Also, although pseudotime is scaled between 0 and 1 and does not correspond to calendar time, it provides a relative measure of amyloid pathology severity and the learned ordering could capture biologically meaningful transitions in early amyloid progression.

Closer examination of the learned progression trajectory revealed amyloid spreading patterns consistent with established findings, with early involvement of the posterior cingulate cortex and precuneus, both core hubs of the default mode network known to be vulnerable to early amyloid accumulation ^12,15^. This agreement with prior observations supports the biological validity of the inferred trajectory and underscores the potential of SLOPE for in vivo disease staging. By modeling progression on a continuous scale, SLOPE facilitates identification of early amyloid changes even in individuals who are globally stable in amyloid burden, offering insights into the sequence of regional amyloid initiation and accumulation.

This study has limitations. The cohort primarily consisted of older adults, limiting characterization of the very earliest stages of amyloid accumulation. Future studies incorporating younger populations may further refine early progression patterns. In addition, SLOPE was applied only to amyloid PET data. Extending the framework to multimodal imaging, including tau PET and MRI, could provide a more comprehensive view of Alzheimer’s disease progression.

## Acknowledgments

Data collection and sharing for the Alzheimer’s Disease Neuroimaging Initiative (ADNI) is funded by the National Institute on Aging (National Institutes of Health Grant U19AG024904). The grantee organization is the Northern California Institute for Research and Education. In the past, ADNI has also received funding from the National Institute of Biomedical Imaging and Bioengineering, the Canadian Institutes of Health Research, and private sector contributions through the Foundation for the National Institutes of Health (FNIH) including generous contributions from the following: AbbVie, Alzheimer’s Association; Alzheimer’s Drug Discovery Foundation; Araclon Biotech; BioClinica, Inc.; Biogen; BristolMyers Squibb Company; CereSpir, Inc.; Cogstate; Eisai Inc.; Elan Pharmaceuticals, Inc.; Eli Lilly and Company; EuroImmun; F. Hoffmann-La Roche Ltd and its affiliated company Genentech, Inc.; Fujirebio; GE Healthcare; IXICO Ltd.; Janssen Alzheimer Immunotherapy Research & Development, LLC.; Johnson & Johnson Pharmaceutical Research & Development LLC.; Lumosity; Lundbeck; Merck & Co., Inc.; Meso Scale Diagnostics, LLC.; NeuroRx Research; Neurotrack Technologies; Novartis Pharmaceuticals Corporation; Pfizer Inc.; Piramal Imaging; Servier; Takeda Pharmaceutical Company; and Transition Therapeutics.

## Conflict of interest

Dr. Saykin has received support from Avid Radiopharmaceuticals, a subsidiary of Eli Lilly (in kind contribution of PET tracer precursor) and holds advisory roles with Siemens Medical Solutions USA, Inc., NIH NHLBI, and Eisai. His editorial commitments include serving as Editor-in-Chief for the journal ‘Brain Imaging and Behavior’, and he participates in various NIH/NIA advisory committees. Dr. Arnold is an inventor on several patent applications on the use of metabolomics in diseases of the central nervous system. He further holds equity in Chymia LLC and IP in PsyProtix and Atai not relevant to this work. Dr. Kaddurah-Daouk is an inventor on a series of patents on use of metabolomics for the diagnosis and treatment of CNS diseases and holds equity in Metabolon Inc., Chymia LLC and PsyProtix. Fahad Mehfooz, Mingzhao Tong, Yipei Wang, Xiaoqian Wang, Shiaofen Fang, Shu Zhang and Jingwen Yan have nothing to disclose.

## Funding Source

This research was supported by NIH grants R01 AG081951, U19 AG074879, U19 AG024904, P30 AG072976, U01 AG068057 and U01 AG072177, as well as NSF grants 2345235 and 1942394.

## Consent Statement

All human subjects provided informed consent and signed consent form.

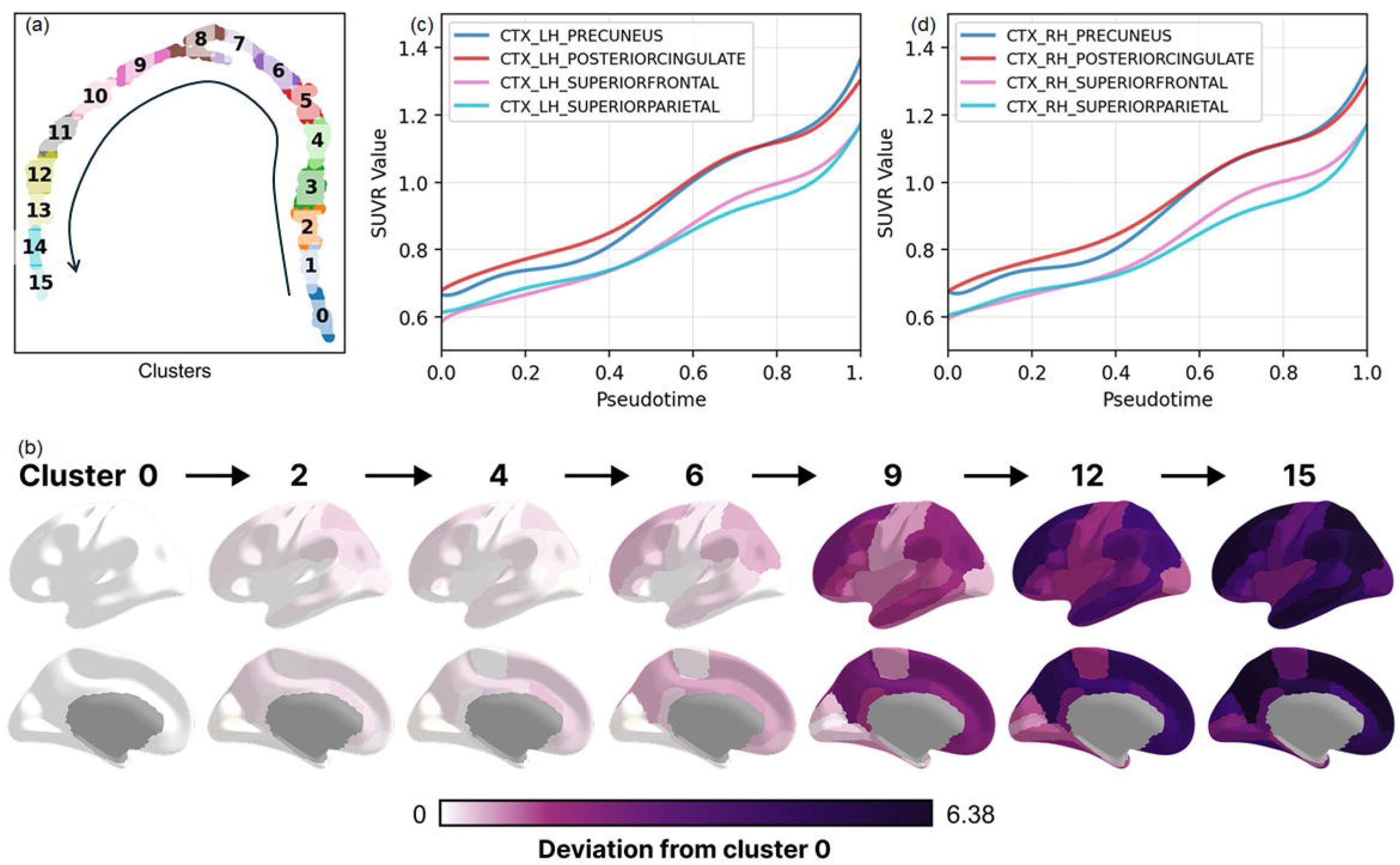

## Notes

### Summary of Updates

Remove markers; Author affiliations updated; Supplemental files updated.

