## Supplementary Text for "Learning a Continuous Progression Trajectory of Amyloid in Alzheimer disease"

**1. Hyperparameter Optimization**

Model hyperparameters were optimized using a hybrid tuning strategy. Grid search was first applied to explore discrete architectural parameters, such as the number of layers, hidden dimensions, and activation functions. This was followed by random search to fine-tune continuous parameters like the learning rate and regularization strengths (i.e., $\lambda_{1}$ and $\lambda_{2}$). For supervised baseline models, 5-fold cross-validation was performed for hyperparameter tuning within the training set to ensure stable performance and mitigate overfitting.

In contrast, SLOPE and the traditional autoencoder are self-supervised models that do not use diagnostic labels during training. These models serve primarily as unsupervised dimensionality reduction techniques, and therefore, the typical goals of cross-validation, such as estimation of prediction performance or supervised parameter tuning, do not directly apply. Like PCA, both SLOPE and autoencoder learn a transformation of the input data exclusively from the training set, and this transformation is subsequently applied to the test set, which remains entirely unseen during model training and hyperparameter tuning. The best SLOPE model was trained for 850 epochs using a five-layer architecture with dimensions [68, 570, 23, 570, 68] and ReLU activation. Optimization was performed with the Adam optimizer and a learning rate of $8.3\times{10}^{-6}$, minimizing mean squared error (MSE) as the loss function. The model balanced its objectives using a reconstruction weight ($\lambda_{1}$) of 0.43 and a directional loss weight ($\lambda_{2})$ of 1.0. In summary, the best SLOPE model generated a 23-dimensional latent embedding for each visit. This embedding was subsequently projected to two dimensions using UMAP for visualization. UMAP was used solely for visualization of the learned latent embeddings and was not part of SLOPE model training. Using the umap-learn package, UMAP was trained in an unsupervised manner on the training-set embeddings and the learned transformation was applied to the test subjects using default parameters without hyperparameter optimization.
